# Extracellular Vesicles-Mediated Crosstalk in Bone: miR-150-5p as a Mechanosensitive Inhibitor of Osteoclastogenesis

**DOI:** 10.1101/2025.09.08.674886

**Authors:** M. Maggio, C. S. Martins, R. Almasri, S. Petrousek, M.Y. Brunet, L.A. Madden, C. Gorgun, T. Ní Néill, C.T. Buckley, L. O’Driscoll, D.A. Hoey

**Author notes:** Corresponding author:* Prof. David Hoey.

## Abstract

Bone is a dynamic tissue that is constantly remodelling via a tightly controlled balance between formation and resorption. However, when osteoclast-mediated resorption exceeds formation, it can lead to net bone loss and the development of osteoporosis. Osteoclastogenesis is regulated by local environmental cues, including paracrine factors released by resident cell populations. Extracellular vesicles (EVs) have recently emerged as key mediators of paracrine communication, though their role in osteoclastogenesis remains underexplored. Therefore, this study investigated how resident mesenchymal-derived bone cells and their secreted EVs modulate osteoclastogenesis, and delineate the signalling factors mediating this anti-catabolic communication. We demonstrate that the secretome of mesenchymal-derived bone cells inhibit osteoclast differentiation to differing degrees depending on the stage of lineage commitment and on the mechanical environment, and demonstrate that this inhibition is mediated via the release of EVs. In addition, we identified the terminally differentiated osteocyte as the optimal parent cell for the production of anti-catabolic EVs and demonstrated the importance EV dosage. Finally, we show that mechanosensitive miR-150-5p is packaged within osteocyte-derived EVs and inhibits osteoclast differentiation. Taken together, this study identifies mechanically-activated osteocyte derived EVs and miR-150-5p as key regulators of osteoclastogenesis and novel molecular therapies for skeletal pathologies.

## Introduction

Bone is a dynamic tissue that has the extraordinary capacity for remodelling and self-renewal to accommodate changes in its biochemical and biophysical environment, maintaining mineral homeostasis and repairing microdamage within the bone matrix. Bone remodelling arises from the coupled action of bone forming osteoblasts and bone resorbing osteoclasts. The balance between osteoblast-driven bone formation and osteoclast-driven bone resorption is tightly controlled, maintaining stable bone mass and mechanical strength under physiological conditions [1]. However, disruption of this balance can result in abnormal remodelling, resulting in bone pathologies such as osteoporosis and osteopetrosis [2, 3]. Osteoporosis arises from an imbalance favouring bone resorption. Driven largely by osteoclasts, this disorder leads to decreased bone mass and mineral density, deterioration of bone microarchitecture and increased bone fragility, resulting in high fracture risk [4]. Osteoclasts are multinucleated bone resorbing cells, and they differentiate from monocyte/macrophage precursors cells upon stimulation with monocyte/macrophage stimulating factor (M-CSF) and receptor activation of NF-κB ligand (RANKL) [5]. While controlled bone resorption is essential for bone remodelling and a healthy skeleton, inhibiting osteoclast activity has been shown to be beneficial in the treatment of osteoporosis [6]. Anti-resorptive treatments, such as bisphosphonates (BPPs), anti-RANKL antibodies (e.g., denosumab), and selective oestrogen receptor modulators (SERMs), function by either inducing osteoclast apoptosis or inhibiting osteoclast recruitment and activity, thereby reducing bone resorption [7]. Although these therapies are effective in maintaining bone density, they have significant limitations, underscoring the need for alternative treatments that can effectively prevent bone loss. Osteoclast formation and function is tightly controlled by biochemical and specifically biophysical cues within the bone microenvironment, often indirectly by signals sent from other mesenchymal derived cells within bone [8]. Therefore, detailing how bone cells regulate osteoclastogenesis and how this may be influenced by their stage of lineage commitment and mechanical environment, is crucial for identifying novel therapeutic strategies for bone disorders.

Bone cells, such as bone marrow stromal/stem cells (MSCs), osteoblasts, and osteocytes have been shown to play a pivotal role in regulating bone remodelling and osteoclast formation. MSCs have been found to have a dual effect on osteoclastogenesis depending on the microenvironment. *In vivo* and *in vitro* studies have demonstrated that MSCs regulation of osteoclastogenesis can occur both via direct contact and paracrine signalling [9, 10]. Under physiological conditions, MSCs can promote or suppress osteoclastogenesis via the secretion of a range of pro- and anti-osteoclastogenic cytokines [11, 12]. In addition, paracrine crosstalk between the more differentiated osteoblast and osteoclast has been proven to be essential to maintain bone homeostasis [13, 14]. Osteocytes terminally differentiate from osteoblasts and become entrapped in the newly formed bone matrix and represent the most abundant cell type in bone [15]. Previous studies have identified osteocytes as master orchestrators of bone remodelling via the control of osteoblast and osteoclast activity [16]. Osteocytes control osteoclastogenesis both via cell-cell interaction and signalling through the release of paracrine factors. While many studies have explored the role of mesenchymal derived bone cells in regulating osteoclastogenesis in isolation, findings can be challenging to directly compare given the differences in cell type and environmental conditions utilised. Therefore, more targeted studies are required to systematically explore how bone cells of the mesenchymal lineage regulate osteoclastogenesis.

A potent regulator of bone remodelling is mechanical loading. Unloading or low load environments have been shown to cause an acceleration of bone turnover, where bone resorbing osteoclasts outpace bone-forming osteoblasts, leading to rapid bone loss. The ability of bone to sense and respond to mechanical stimuli is orchestrated by the cells of the osteogenic lineage, the MSCs [17], the osteoblast [18], and particularly the osteocyte [19]. These cells are subjected to a constantly changing mechanical environment that comprises various biophysical stimuli, such as strain, stress, shear, pressure and fluid flow, which they convert into biochemical signals driving cellular responses. As mentioned above, mesenchymal-derived cells regulate osteoclastogenesis mainly via paracrine signalling. Secretion of signalling molecules can change in response to a wide variety of stimuli, including mechanical stimulation [20]. It is widely accepted that the osteocytes are the primary mechanosensors in bone and regulate bone resorption in response to mechanical loading [21–23]. Interestingly, recent *in vivo* studies have also demonstrated the mechanosensitive nature of both MSCs and osteoblasts, highlighting their contribution to mechanically mediated bone remodelling [24]. Consistent with these findings, *in vitro* studies have demonstrated that mechanical stimulation prompts MSCs to alter their secretome suppressing osteoclast formation thereby contributing to rebalance bone remodelling [20]. Similarly, mechanical loading has been shown to modulate osteoblast and osteocyte secretome, resulting in a mechanically driven osteoclast inhibition [25–28]. These mechanically induced changes in the bone cell regulation of osteoclastogenesis warrant further investigation of the specific mechanically regulated signalling components mediating these effects, as they may represent novel anti-catabolic therapies.

Extracellular Vesicles (EVs) have been widely recognised as important mediators of intercellular communication [29]. EVs are lipid bilayer membrane-bound particles released by cells into the extracellular space both in physiologic and pathologic conditions. Over the last few decades, numerous studies have demonstrated that EVs carry and transfer a cargo of biomolecules including proteins, mRNA, and miRNA to regulate pathways in the target cell [30]. In particular, there is a growing body of evidence indicating that bone-derived EVs miRNAs can interact with different signalling molecules to regulate the functions of osteoblasts and osteoclasts, thereby influencing the bone remodelling process [31, 32]. The presence of specific groups of biomolecules in EVs suggests the existence of a sorting mechanism that coordinates the selective packaging of RNAs and proteins. Biophysical cues and biochemical cues have been used to tune EVs cargo and enhance their therapeutic potential [33, 34]. For instance, EVs secreted by bone cells have been shown to have a pivotal role in coordinating bone remodelling and our group has previously demonstrated that EVs secreted by mechanically stimulated bone cells can drive an anabolic response by regulating MSCs recruitment, osteogenesis and vessel formation. Additionally, our findings indicate that mechanical loading influences the miRNA content of osteocyte-derived EVs, pointing to a potential miRNA-mediated mechanism driving regenerative effects [35, 36]. However, the role of bone cell secreted EVs and associated miRNAs in regulating osteoclastogenesis has yet to be fully explored to date.

Therefore, the aim of this study is to investigate the role of mesenchymal-derived bone cells and secreted EVs in the coordination of osteoclastogenesis and to determine how the stage of lineage commitment and mechanical environment of the parent cell influences this process. Moreover, we aim to identify the signalling components within bone cell-derived EVs that may mediate the effects on osteoclastogenesis, offering new avenues for therapeutic intervention.

## Methods

### Cell culture

#### Mesenchymal stromal/stem cell culture

Human mesenchymal stem cells (MSCs) were isolated from bone marrow (Lonza) and cultured in Dulbecco’s modified eagle medium (DMEM) with GlutaMAX™ (Gibco) supplemented with 10 % Fetal Bovine Serum (FBS) (Gibco) and 1 % Penicillin-Streptomycin (P-S) (Sigma). Cells were cultured at 37 °C and 5 % CO_2_ and used between passage 3-5 were for all the experiments, as previously described [35, 37].

#### Osteoblast culture and differentiation

MSCs were seeded at 6,500 cells/cm2 in a 6 well-plate (Sarstedt) and cultured in DMEM GlutaMAX™ (Gibco) supplemented with 10 % FBS and 1 % P-S at 37 °C and 5% CO_2_. 100 nM dexamethasone, 0.05 nM L-ascorbic acid and 10 mM β-glycerol phosphate were added to the media to induce osteogenic differentiation and were cultured for further 14 days to obtain osteoblasts (OB). At the endpoint, OB were washed in PBS, then 1 mL/well of 2 mg/ml collagenase type II in PBS (Gibco) was added for 30 minutes to disrupt the deposited matrix. Collagenase was then removed and 1 mL/well of trypsin (Sigma) was added to lift the OBs for reseeding onto glass slides prior mechanical stimulation.

#### Osteocyte culture

As human osteocytes are challenging to obtain in sufficient number and purity, the murine osteocyte-like MLO-Y4 cell line (Kerafast) was chosen for this study [38]. MLO-Y4 cells maintained in MEM Alpha (α-MEM) (Merk) supplemented with 2.5 % FBS (Gibco), 2.5% Calf Serum (CS) (Biosera), 1 % P-S (Sigma), and 1 % L-Glutamine (L-G) (Sigma) as previously described [39] . Cells were cultured at 37 °C and 5 % CO_2_ and used for experiments when they had reached 80 % confluency between passages 40-60.

### Mechanical stimulation and conditioned media collection

Glass slides (75x38 mm) (Corning) were sterilised in 70 % IMS for 1 h, allowed to dry, and coated with rat tail collagen type I (0.15 mg/mL) (Corning) for 1 h. Prior to seeding a wash step was performed with PBS (Sigma). MLO-Y4 cells were seeded with a density of 11,600 cells/cm^2^ onto the coated glass slides. MSCs and OB were seeded independently on the coated glass slides at a density of 6,140 cells/cm^2^. After 24 h culture, cells were treated with serum reduced media overnight. Osteocyte (OCY) media consisted of α-MEM supplemented with 0.5 % FBS, 0.5 % CS, 1 % L-G, 1 % P-S, while the MSCs and OB media consisted of DMEM GlutaMAX™ supplemented with 0.5 % FBS, 1 % P-S. The slides were then placed in a custom-made parallel plate flow chamber (PPFC) bioreactor as previously described [40]. Briefly, the PPFCs were connected to a syringe pump to mechanically stimulate the cells by applying an oscillatory fluid flow included shear stress of 1 Pa at 1 Hz for 2 h. Cells were also cultured statically in the PPFCs for 2 h as a control. Following stimulation, the chambers were disassembled, the slides were washed with PBS and put in custom made slide chambers with 2.5 mL of α-MEM supplemented with 0.5 % EVs-depleted FBS (dFBS), 0.5 % EVs-depleted CS (dCS), 1 % L-G and 1 % P-S for osteocytes and DMEM supplemented with 0.5 % dFBS and 1 % P-S for MSCs and OB. After 24 h of culture, static conditioned media (CM-S) and fluid shear stimulated conditioned media (CM-F) were collected and centrifuged at 3,000 g for 10 min at 4 °C to remove debris. Supernatant was collected and stored at - 80 °C.

### Bone cell regulation of osteoclastogenesis

The use of human blood samples for this study was approved by the research ethics committee of Trinity College Dublin and was conducted in accordance with the Declaration of Helsinki. Leukocyte-enriched buffy coats from anonymous healthy donors were obtained with permission from the Irish Blood Transfusion Service (IBTS), St. James’s Hospital, Dublin (Researcher code: 0000RES625). Donors provided informed written consent to the IBTS for their blood to be used for research purposes. Peripheral blood mononuclear cells (PBMCs) were isolated from leukocyte-enriched human buffy coats using density gradient centrifugation with LymphoPrepTM solution (Progen). CD14^+^ monocytes were isolated from PBMCs by magnetic sorting using MagniSort Human CD14 Positive Selection Kit (Invitrogen) according to the manufacturer’s instructions. CD14^+^ monocytes were seeded at 625,000 cells/cm^2^ in 96-well plates in α -MEM containing 10 % FBS, 1 % P-S and 1 % L-G supplemented with 25 ng/mL M-CSF (Myltenyi Biotec) and 50 ng/mL Recombinant Human sRANK Ligand (RANKL) (PeproTech) (+RANKL) to induce osteoclastogenesis for 7 days. Groups cultured in growth media supplemented with 25 ng/mL M-CSF acted as a negative control for osteoclastogenesis (-RANKL). To assess the effect of bone cell secretome on osteoclastogenesis, CM collected from statically or mechanically stimulated cells was supplemented with 9 % FBS and added in a ratio of 1:1 along with growth media in presence of 50 ng/mL RANKL and 25 ng/mL M-CSF. Media and CM were replenished every 3.5 days. At day 7 cells were fixed with 2 % paraformaldehyde (PFA) for 15 min at room temperature, then washed twice with PBS and stained for tartrate-resistant acid phosphatase (TRAP) activity with a leukocyte acid Phosphatase (TRAP) Kit (Sigma) according to the manufacturer’s protocols. Finally, cells were incubated with Triton X-100 0.1 % (Sigma) for 10 min, rinsed with PBS and then incubated with DAPI (1:2000) (Sigma) for 10 min. TRAP-positive multinuclear cells containing 3 or more nuclei were counted as osteoclasts. Cells were imaged using an Olympus IX83 inverted microscope with 10x objective and osteoclasts, number of nuclei per osteoclast, and total number of cells were manually counted using the cell counter plugin on Image-J software (version 1.54p, NIH, USA).

### Extracellular vesicles collection

EVs were collected from CM via ultracentrifugation using a 70Ti-fixed angle rotor as previously described [36]. Briefly, CM was centrifuged at 2,000 g for 15 min at 4 °C and then filtered through a 0.45 µm pore filter. Next, CM was ultracentrifuged at 110,000 g for 75 min at 4 °C. The resulting EVs pellets were washed in PBS and ultracentrifuged again at 110,000 g for 75 min at 4 °C. Finally, EVs pellets were resuspended in PBS and stored at -80 °C.

### Extracellular vesicles characterisation

#### Bicinchoninic acid (BCA) assay

EVs sample of 50 µL was suspended in 50 µL of cell lysis buffer (Invitrogen) and 25X proteinase inhibitor (Roche). EVs lysate was incubated on ice for 30 min, vortexing every 10 mins for 10 s. After 30 min, the sample was centrifuged at 13200 rpm at 4 °C for 10 min. The supernatant was transferred to a new tube and stored at -20 °C until required. EVs lysate was quantified using the Bio-Rad protein assay dye reagent (Bio-Rad). Bovine serum albumin (BSA) standards of 12.5-800 µg/mL were prepared in PBS and used to calculate the protein content of the samples. Lysed EVs was diluted 1:2 in PBS and whole cell lysate were diluted 1:20 in PBS. 10 µL of standards and sample was added to a 96-well plate in duplicate. Bio-Rad dye reagent was diluted 1:5 in deionised water and 200 µL added to each standard and sample. Absorbance was read at 570 nm using a fluoStar optima microplate reader (Serial #: 08-100-241) and protein levels were calculated from the standard curve.

### Flow Cytometry

EVs surface antigens were exposed to antibodies diluted in 0.22 μm-filtered PBS with 2 % dFBS supplemented with protease inhibitor and phosphatase inhibitor (IFCM buffer). The antibodies used were anti-CD63 conjugated with FITC (1:150) (Biolegend), CD9-PE (1:5000) (Biolegend), CD81-PE-Cy7 (1:150) (Biolegend). The EVs were incubated with the antibodies for 45 min at room temperature in the dark, and washed using a 300kDa filter (Nanosep), resuspended in 50 μL IFCM buffer and acquired within 2 h on the ImageStream X MK II imaging flow cytometer (Amnis/Luminex, Seattle, USA) at 60x magnification and low flow rate. EVs-free IFCM buffer, unstained EVs, single-stained controls and fluorescence minus one (FMO) controls were run in parallel. Fluorescence was within detection linear range in the following channels: FITC was measured in channel 2 (B/YG_480-560 nm), PE in channel 3 (B/YG_560-595 nm), PE-Cy7 in channel 6 (B/YG_ 745-780 nm) and APC in channel 11 (R/V_642-745 nm). Brightfield in channel 1 and 9 (B/YG_435-480 and R/V_560-595 nm filter, respectively) and side scatter channel (SSC) in channel 12 (R/V_745-780 nm 49 filter). Data analysis performed using IDEAS software v6.2 (Amnis/Luminex, Seattle, USA). EVs were gated as SCC-low vs fluorescence, then as non-detectable brightfield (Fluorescence vs Raw Max Pixel Brightfield channel), gated EVs were confirmed in IDEAS Image Gallery.

### Nanoparticle tracking analysis

Nanoparticle tracking analysis was performed on EVs suspensions using the NTA NS500 system (NanoSight, Amesbury, UK) to determine particle size based on Brownian motion. EVs samples were diluted at 1:50 in PBS and injected into the NTA system, which obtained four 40-second videos of the particles in motion. Videos were then analysed with the NTA software to determine particle size.

### Transmission electron microscopy

A 20 µL sample of EVs suspension was placed onto parafilm (Sigma-Aldrich). A formvar carbon-coated nickel grid (Ted Pella Inc) was placed on top (coated side facing the droplet) of the EVs suspension droplet. The grid was incubated for 60 min at room temperature, washed in 30 µL of PBS (x3 times) on parafilm for 5 min. Absorbent paper was used to remove excess PBS from the washing steps. A droplet of PFA (2 %) was placed on parafilm, and the grid was placed on top and fixed for 10 min. The PBS washing steps were repeated. The grid was then contrasted in 2 % uranyl acetate (BDH) and all images were taken using the JEOL JEM-2100 transmission electron microscope at 120 kV.

### Super-resolution dSTORM microscopy

Super-resolution fluorescent microscopy analyses of EVs were performed with 100x oil immersion 1.45 NA objective using Nanoimager S Mark II microscope from ONI (Oxford Nanoimaging, Oxford, UK) equipped with 405 nm/150 mW, 488 nm/1 W, 560 nm/500 mW, 640 nm/1 W lasers. Fluorescence excited by 488, 560 and 640 lasers was recorded using band pass filters 498-551 nm, 576 - 620 and 665-705 nm respectively. Fluorescence excited by 488, 560 and 640 lasers was recorded using band pass filters 498-551nm, 576 - 620 and 665-705nm respectively. ONI EV Profiler Kit 2 (EVP2) were used to prepare EVs samples for dSTORM as per manufacturer’s instruction. This kit includes panEV stain (membrane dye), tetraspanin antibodies (CD9, CD63, CD81), capture chips, capture reagents and ONI BCubed dSTORM Imaging Buffer used in this study. Three-channel (647, 560 and 488) dSTORM data (1000 frames per channel) were acquired sequentially using AutoEV CODI acquisition software using 180 mW power at the objective for 488 nm laser, 45mW for 560 nm laser and 170mW for 640 nm laser at 40 ms exposure per frame. For panEV stain 3,000 images were acquired. Before each imaging session, beads slide calibration was performed to align fluorescent channels, achieving a channel mapping precision smaller than 10 nm. Single-molecule localization, clustering and positivity counting was performed using default EVP2 app setting within the CODI software provided by ONI (https://alto.codi.bio). Data was exported as report and csv file for further analysis.

### Extracellular vesicles regulation of osteoclastogenesis

To determine whether EVs released by mesenchymal derived bone cells influence osteoclastogenesis, human blood derived CD14^+^ monocytes were seeded at 625,000 cells/cm^2^ in 96-well plates in α -MEM containing 10 % dFBS, 1 % P-S and 1 % L-G (growth media) supplemented either with 25 ng/mL M-CSF (Myltenyi Biotec) and 50 ng/mL RANKL (PeproTech) (+RANKL) to induce osteoclastogenesis for 7 days or with 25 ng/mL M-CSF alone (-RANKL) as a control. EVs enriched media was prepared by supplementing growth media with 0.5 µg/mL of MSCs, osteoblast and osteocyte derived static EVs (EV-S) and fluid shear stimulated EVs (EV-F) in the presence of M-CSF and RANKL. To assess the effect of secreted soluble factors not packaged within EVs, CM-S and CM-F depleted of EVs (DM-S and DM-F) were supplemented with EVs-depleted FBS and added in a ratio of 1:1 along with growth media in presence of both M-CSF and RANKL. Media and EVs were replenished after 3.5 days. Osteoclast differentiation was assessed by TRAP and DAPI staining as described above. Cells were imaged using Olympus IX83 inverted microscope with 10x objective and osteoclasts, number of nuclei per osteoclast and total number of cells were manually counted using the cell counter plugin on Image-J software (version 1.54p, NIH, USA).

To assess the influence of EVs concentration on osteoclastogenesis, EVs enriched media was prepared by adding 0.5, 1, 2, and 4 µg/mL of both static and fluid shear stimulated osteocyte derived EVs to the growth media. Human blood derived CD14^+^ were cultured in EVs enriched media in a ratio of 1:1 along with growth media in the presence of M-CSF and RANKL for 7 days as described above. Osteoclast were stained for TRAP and DAPI staining and imaged using Olympus IX83 inverted microscope with 10x objective and quantification was performed using Image-J software (Version 1.54p, NIH, USA).

### Osteoclast resorption activity assay

Osteoclast activity was assessed by determining the area of resorption on bovine bone slices. Briefly, bone slices (6mm diameter, 200-220 µm thickness) were cut from bovine cortical femur with a low-speed water-cooled diamond saw (Beuhler). 24 hours before seeding, bone slices were sterilized in 70 % ethanol, dried and incubated in media overnight. Subsequently, CD14+ monocytes were cultured on the sterile bone slices in a 96-well plate at a density of 200,000 cells per bone slice in αMEM supplemented with 10 % dFBS, 1 % P-S and 1 % L-G, in the presence of M-CSF (25 ng/mL) and RANKL (50 ng/mL) for 14 days. The effect of osteocyte derived EVs on osteoclast resorption activity was assessed by adding EVs enriched media, containing 0.5 µg/mL EV-S and EV-F in a ratio of 1:1 with growth media, in the presence of M-CSF and RANKL. Media and EVs were replenished every 3.5 days. After culture, samples were washed with PBS and 200 µL of deionised water was added for 30 min. Subsequently, samples surface was scraped with a cotton swab to remove cells. Finally, samples were serially dehydrated in an increasing concentration of ethanol and a final drying step in hexamethyldisilazane to remove all water before being imaged by scanning electron microscopy (SEM). After dehydration, samples were left to dry on adhesive double-sided carbon tape attached to 11.7 mm diameter aluminium stubs (Micro to Nano) overnight. Samples were then coated with a 5 nm thick coating of gold/palladium using a sputter coater and imaged in a Zeiss Ultra SEM (Zeiss). Representative images were taken using a secondary electron detector with an accelerating voltage of 10 kV. Images for quantitative analysis were taken using an in-lens detector under identical conditions. Resorption pit area was calculated using Image-J software (Version 1.54p, NIH, USA).

### Bioinformatic analysis

Building on our previous findings highlighting that mechanical stimulation of osteocytes led to the productions of EVs significantly more enriched in miR-150-5p compared to those from static condition [36], miR-150-5p human target genes were identified using TargetScan (version 8.0, https://www.targetscan.org/vert_80/). Functional annotation and enrichment analysis of the selected genes were performed using the Database for Annotation, Visualization, and Integrated Discovery (DAVID) [41], focusing on Gene Ontology Biological Process (GO BP) terms. The threshold for significance was set to p-value ≤ 0.05 (EASE score) and Enriched GO BP terms were visualized through a bubble plot generated in R Studio (version 2025.05.1, Build 513) using the ggplot2 package.

### Human monocytes transfection and osteoclastogenesis assessment

For CD14^+^ monocyte transfection, cells were seeded 625,000 cells/cm^2^ in 96-well plates in α -MEM containing 10 % FBS, 1 % P-S and 1 % L-G supplemented with 25 ng/mL M-CSF (Myltenyi Biotec) and 50 ng/mL RANKL (PeproTech) and cultured for 3.5 days prior to transfection. miRNA complexes were formed through electrostatic interaction of RALA (GenScript) [42] at an N:P ratio of 10 [43] for miRNA mimic hsa-miR-150-5p and Mimic Negative Control #1 (Scrambled control) (Horizon Discovery) at concentrations of 1 ng/µL and 5 ng/µL in OptiMEM (Gibco). Complexes were added to the cells following PBS wash and incubated for 5h at 37°C. + RANKL positive control was incubated with OptiMEM only in for 5 h at 37 °C. At the end of the incubation, complexes were removed, and cells were cultured in the same media formulation as before transfection for additional 3.5 days. At the end of culture, osteoclast differentiation was assessed by TRAP and DAPI staining as described above. Cells were imaged using Olympus IX83 inverted microscope with 10x objective and osteoclasts, number of nuclei per osteoclast and total number of cells were manually counted using the cell counter plugin on Image-J software (version 1.54p, NIH, USA).

### Statistical analysis

All data were analysed using GraphPad Prism 10.2.2. All data were analysed using an ordinary one-way or two-way ANOVA, assuming Gaussian distribution. Statistically significant differences were indicated as *p<0.05, **p<0.01, ***p<0.001, **** p<0.0001.

## Results

### Mesenchymal-derived bone cells secrete paracrine factors that regulate osteoclastogenesis in a manner that is dependent on the stage of lineage commitment and the mechanical environment

To investigate the extent to which cells of the mesenchymal lineage can regulate osteoclastogenesis, osteoclast differentiation in response to conditioned media (CM) collected from stem/stromal cells, osteoblasts, and osteocytes cultured statically (CM-S) or exposed to 2 h oscillatory fluid shear (CM-F) was evaluated.

Human blood derived CD14^+^ monocytes cultured for 7 days in the presence of M-CSF and RANKL resulted in the formation of TRAP-positive multinucleated cells, demonstrating the ability of this cell type to differentiate into mature osteoclasts. Treatment with CM collected from MSCs cultured in static or mechanically stimulated conditions resulted in a significant inhibition of osteoclast formation. Specifically, the secretome from statically cultured MSCs (CM-S) demonstrated a robust near total inhibition of osteoclastogenesis (99 % reduction, p<0.0001) when compared to +RANKL controls. Interestingly, the application of fluid shear to the MSCs (CM-F) altered the secretome resulting in a less inhibitory response (35 % reduction compared to +RANKL, p<0.0001), (Error! Reference source not found. A, B). Cell nuclei number was also assessed as a proxy for osteoclast maturity and quantification demonstrated that osteoclast maturation was also reduced following treatment with MSCs CM when compared to +RANKL control (Error! Reference source not found. A, C). A key step in the formation of osteoclasts is the adherence of the CD14+ monocyte to the surface [44]. It was visually evident (Figure 1 A) that factors released from statically cultured MSCs reduced the adhesion of monocytes to the culture plate, which was further validated upon quantification (Figure 1 A, D). This reduction in adhesion was not seen following treatment with CM-F. Taken together, this data indicates that statically cultured MSCs significantly inhibit osteoclastogenesis in part via the secretion of factors that reduces monocyte adhesion, an effect that is diminished following mechanical stimulation.

**Figure 1.**
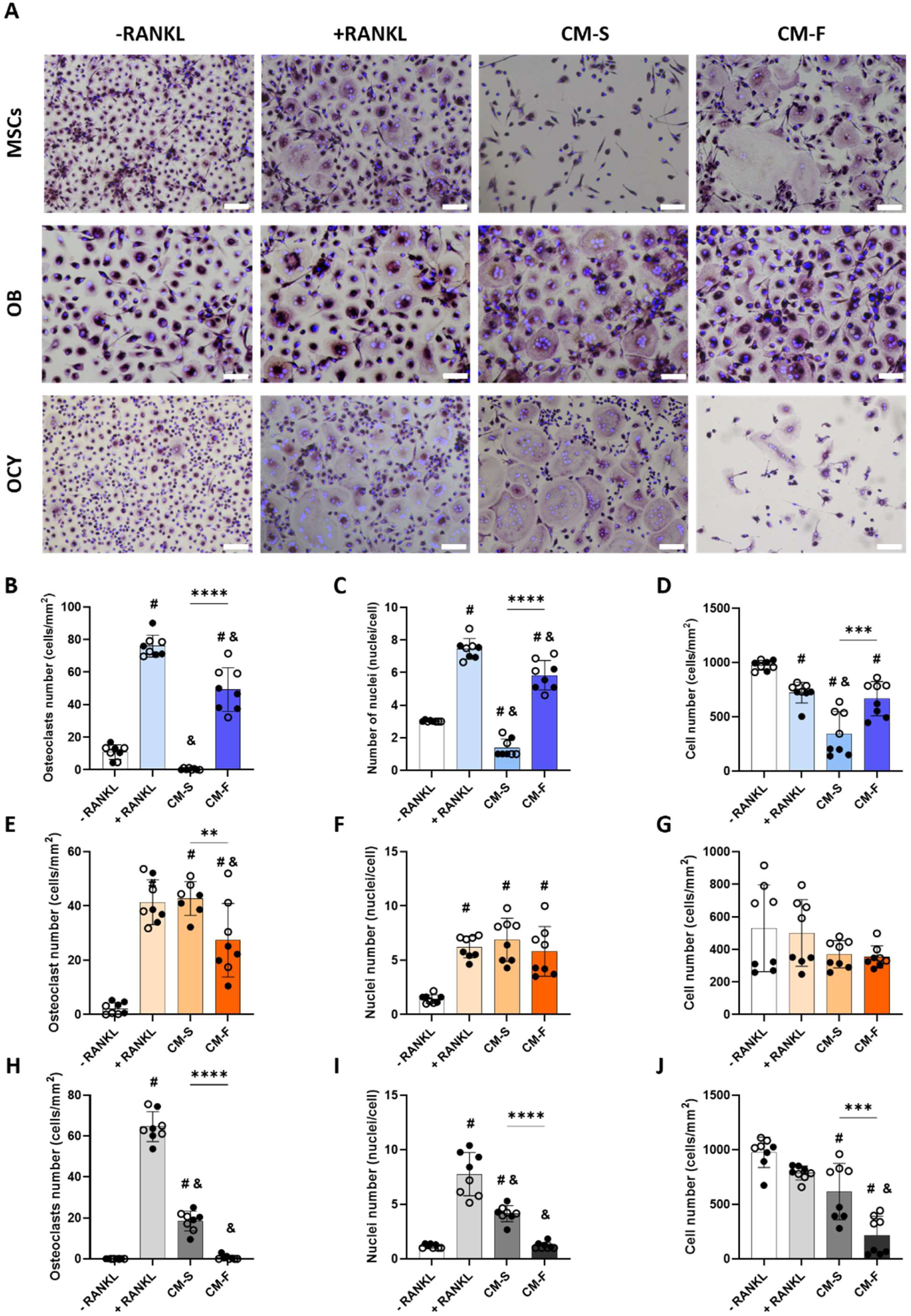
Bone cells from the mesenchymal lineage secrete paracrine factors that inhibit osteoclastogenesis in a manner that is dependent on the stage of lineage commitment and mechanical environment. (A) TRAP and DAPI staining of human monocytes following 7 days treatment with M-CSF (-RANKL); M-CSF and RANKL (+RANKL); M-CSF, RANKL and static MSCs, OB or OCY conditioned media (CM-S); and M-CSF, RANKL and mechanically activated MSCs, OB or OCY conditioned media (CM-F). (B, E, H) Quantification of osteoclast number (identified as TRAP+ cells with ≥3 nuclei), (C, F, I) nuclei number and (D, G, J) total cell number for MSCs, OB, and OCY CM groups respectively. Scale bar 100 µm. Values are mean ± SD for 2 different human donors (● = Donor 1, ○ = Donor 2). *p<0.05, **p<0.01, ***p<0.001, **** p<0.0001, #p < 0.01 vs -RANKL, & p< 0.0001 vs +RANKL.

To assess the effect of mesenchymal lineage commitment on the regulation of osteoclastogenesis, MSCs were differentiated into osteoblasts for 14 days (OB) before CM was collected from both statically and mechanically stimulated cultures [45]. Unlike the parent MSCs, the secretome from statically cultured OB does not contain paracrine factors that inhibit osteoclastogenesis. However, following mechanical stimulation of the osteoblast, CM treatment results in a significant inhibition of osteoclastogenesis when compared to the static group and RANKL positive control (34 %, p< 0.01, and 36 %, p<0.05, reduction in osteoclastogenesis respectively) (Figure 1 A, E). No differences in osteoclast maturity were observed between the + RANKL control and the CM treated groups (Figure 1 A, F). This inhibitory effect of the mechanically stimulated OB secretome was not due to effects on monocyte adhesion as no difference was identified between treated groups (Figure 1 A, G). Therefore, the commitment of MSCs to the osteogenic lineage alters the osteoclastogenic properties of the secretome.

Osteocytes represent the terminally differentiated state of the mesenchymal lineage and their role in coordinating osteoclastogenesis is well studied [46]. Both CM collected from osteocytes cultured in static or mechanically stimulated conditions resulted in a significant inhibition of osteoclast formation. The secretome from statically cultured osteocytes elicited a 72 % reduction (p<0.0001) in osteoclast formation which was further augmented following mechanical stimulation (99 % reduction, p<0.0001) when compared to +RANKL controls (Figure 1 A, H). This enhanced anti-catabolic property following mechanical stimulation is consistent with that seen in OB. Similarly, nuclei number quantification identified a decreased number of nuclei upon treatment with CM-S which was further reduced with CM-F, indicating a lower level of osteoclast maturation (Figure 1 A, I). Interestingly, monocyte adhesion was not affected by treatment with CM collected from static osteocytes. However, treatment with CM collect following osteocyte mechanical stimulation resulted in a significant decrease in monocyte adhesion (Figure 1 A, J). This data is consistent with the previous identified role of the osteocyte in the regulation of osteoclasts and indicates that mechanically stimulated osteocyte may inhibit osteoclastogenesis in part via the secretion of factors that reduces monocyte adhesion. Taken together these data demonstrate that mesenchymal derived bone cells can regulate osteoclastogenesis and maturation via a paracrine mechanism that is strongly dependent on the stage of lineage commitment and the mechanical environment at each stage of the cell lineage.

### Mesenchymal derived bone cells secrete extracellular vesicles that are of similar size but differ in quantity and surface marker expression at each stage of lineage commitment

EVs were collected and characterised from media conditioned by MSCs (MSCs-EVs), osteoblasts (OB-EVs) and osteocytes (OCY-EVs) as per MISEV guidelines [47]. NTA analysis was performed to determine their respective size distribution and concentration. As displayed in Figure 2 A and B, similar size distribution profiles were observed between all EVs groups ranging from 100 to 200 nm with respective average diameters of 163 nm, 160 nm, and 160 nm for MSCs-EVs, OB-EVs, and OCY-EVs. Interestingly, normalising the EVs concentration to the parent cell seeding density, highlighted a significantly greater EVs average yield from osteoblasts (6.99 x 10^9^ NPS/mL/10^6^ cells) compared to EVs secreted from MSCs (2.64 x 10^9^ NPS/mL/10^6^ cells) or osteocytes (2.23 x 10^9^ NPS/mL/10^6^ cells) (Figure 2 C). The morphology of these vesicles was evaluated by TEM revealing nanosized particles (<200 nm) with a spherical shape (white arrow) for all three groups which is consistent with the NTA analysis (Figure 2 F). The total protein content in EVs lysates was determined via a BCA assay and exhibited a similar trend in protein level to that seen in the nanoparticle concentration where there is a slight but non-significant increase in protein quantity in the OB-EVs group when compared to the MSCs and OCY-EVs (Figure 2 E). Finally, the presence of tetraspanin markers (CD9, CD63 and CD81) was assessed by flow cytometry and super resolution microscopy. All three groups were found positive for the three markers with no significant variations as the cells progress through the lineage (Figure 2 D, G).

**Figure 2.**
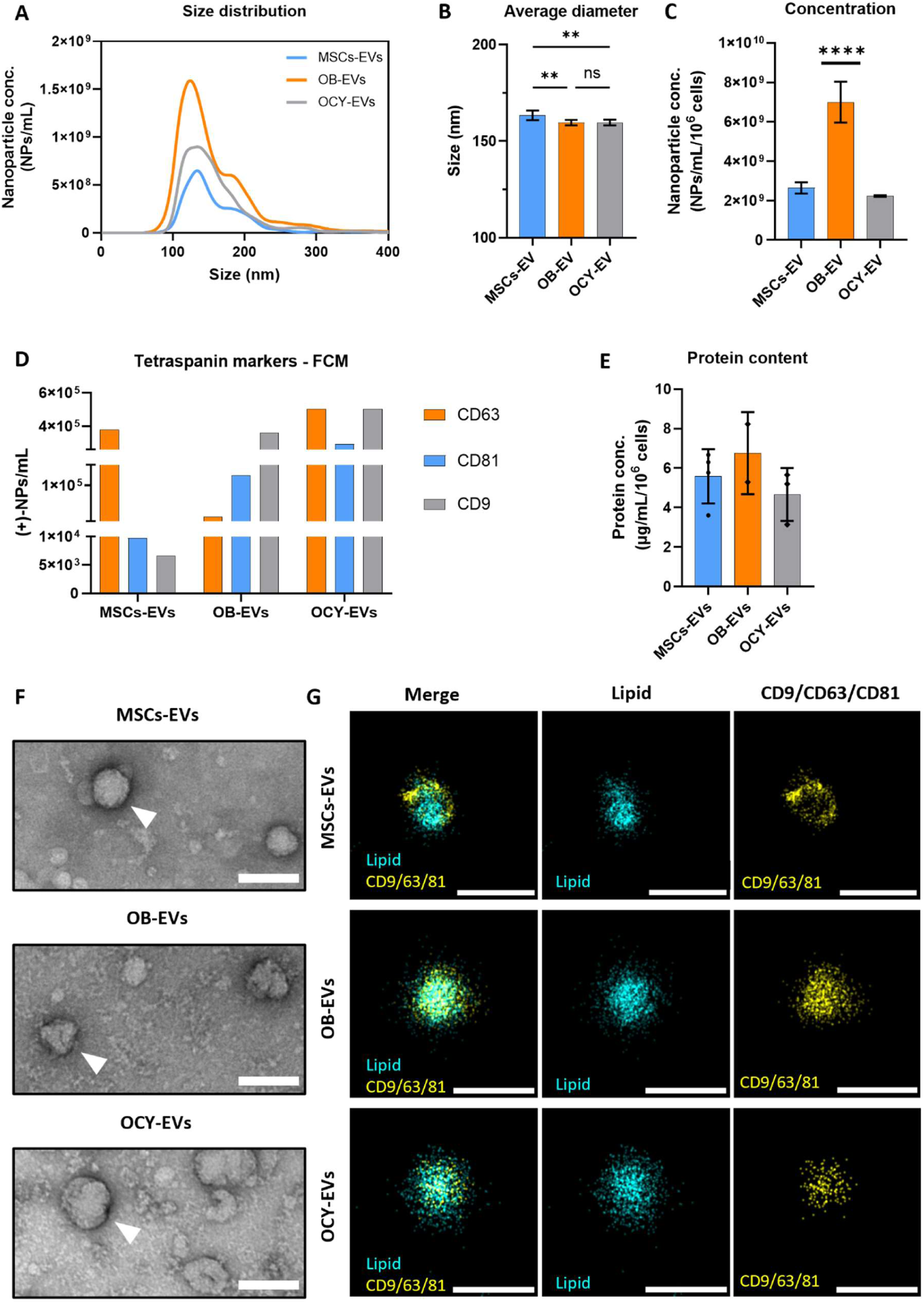
Comparative isolation and characterisation of EVs derived from the conditioned media of mesenchymal stem cells (MSCs), osteoblasts (OB) and osteocytes (OCY). Nanoparticle Tracking Analysis (NTA) was performed to determine (A) Size Distribution (B) Average diameters and (C) Nanoparticle Concentrations. (D) Detection of tetraspanin markers CD63, CD81 and CD9 via flow cytometry. (E) Protein content via the bicinchoninic acid assay. (F) TEM observations with scale 100nm (white arrow indicating EVs). (G) Super resolution microscopy observations of single nanoparticle with the fluorescence detection of membrane (lipid stain) and trio-tetraspanin markers (CD9/CD63/CD81) with scale bar = 200 nm. Statistical analysis performed via one-way ANOVA with post-hoc Tukey test, P values displayed as follow p*≤ 0.05, p**≤ 0.005, ***p≤ 0.001, **** p≤0.0001.

### Mesenchymal derived bone cells secrete extracellular vesicles that inhibit osteoclast differentiation

Given the key role that extracellular vesicles (EVs) play in coordinating cell-cell communication, we next sought to investigate the role that EVs may play in the above demonstrated paracrine signaling between mesenchymal-derived bone cells and osteoclasts under both static conditions (EV-S) or following mechanical stimulation (EV-F). The EVs dose used was 0.5 µg/mL, equivalent to the average EVs protein concentration found in the CM from each cell type. In addition to collected EVs, the effect of static and mechanically activated CM that was depleted of EVs (DM) was also investigated to assess whether soluble factors within the CM, not packaged within EVs, may influence osteoclast differentiation.

EVs released by statically cultured and mechanically stimulated MSCs significantly inhibited osteoclastogenesis, independent of the mechanical environment. Similarly, CM collected from statically cultured and mechanically stimulated MSCs that was depleted of EVs also demonstrated an inhibition of osteoclastogenesis, indicating that soluble factors not packaged within EVs can also affect this process (Figure 3 A, B). Furthermore, quantification of osteoclast nuclei showed that osteoclasts differentiated upon treatment with EVs or CM depleted from EVs (DM) displayed decreased number of nuclei per cell (Figure 3 A, C), which is consistent with what was identified with full conditioned media treatment. In addition, monocyte adhesion was not found to be impaired by EVs or soluble factors contained within the DM independent of mechanical stimulation of the MSCs (Figure 3 A, D).

**Figure 3.**
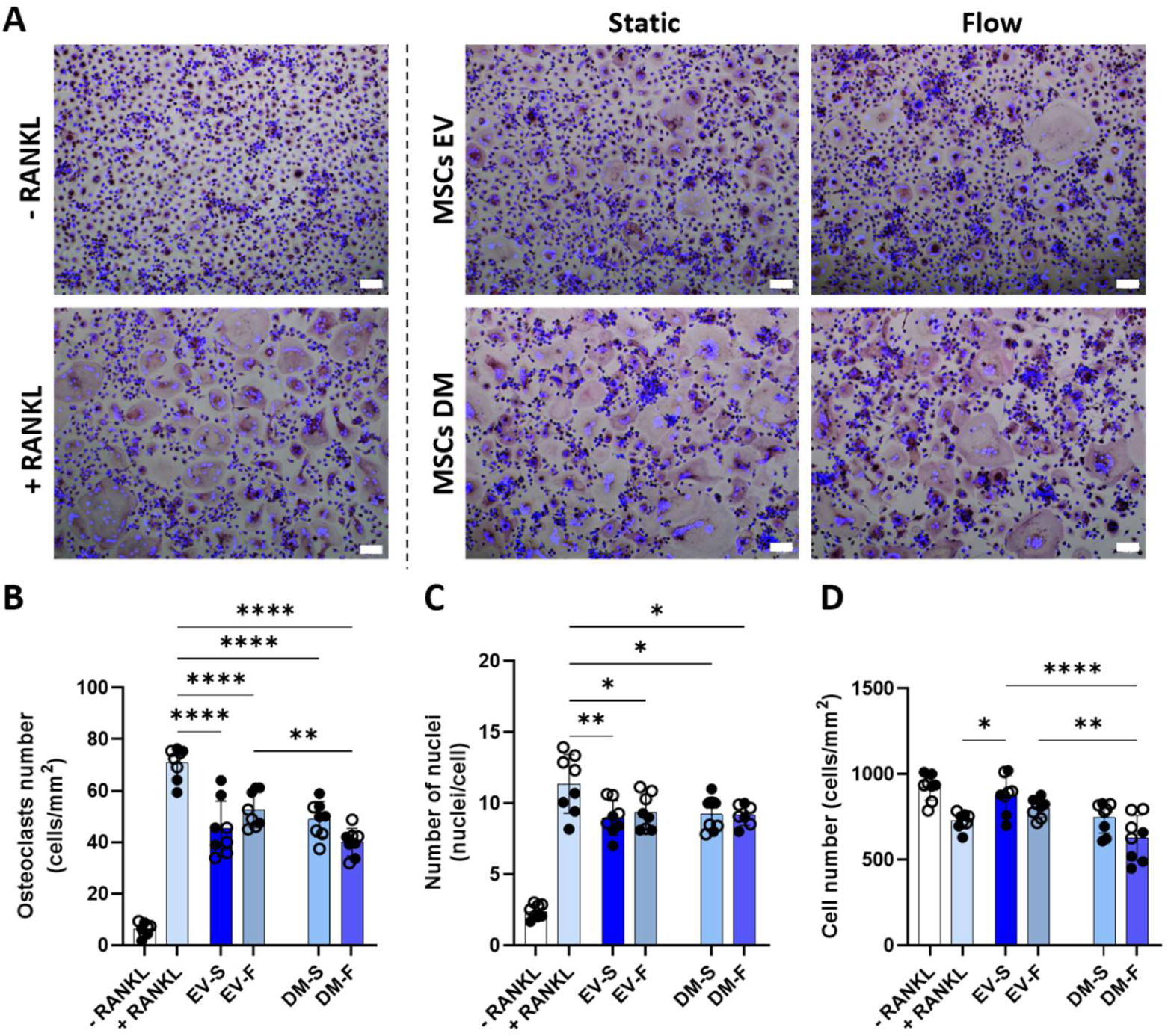
MSCs secrete extracellular vesicles that inhibit osteoclastogenesis independent of mechanical stimulation. **(A)** TRAP and DAPI staining of human monocytes following 7days treatment with M-CSF (-RANKL); M-CSF and RANKL (+RANKL); M-CSF, RANKL and 0.5 µg/mL static MSCs derived EVs (EV-S); M-CSF, RANKL and 0.5 µg/mL mechanically activated MSCs derived EVs (EV-F); M-CSF, RANKL and static EVs depleted MSCs CM (DM-S); and M-CSF, RANKL and mechanically activated EVs depleted MSCs CM (DM-F). **(B)** Quantification of osteoclast number (identified as TRAP+ cells with ≥3 nuclei), **(C)** nuclei number and **(D)** total cell number. Scale bar 100 µm. Values are mean ± SD for 2 different human donors (● = Donor 1, ○ = Donor 2). *p<0.05, **p<0.01, ***p<0.001, **** p<0.0001.

Moving forward along the mesenchymal lineage, EVs secreted by osteoblasts both in static culture and following mechanical stimulation were shown to possess anticatabolic potential. Specifically, osteoblasts cultured in static conditions produced EVs that led to a 25 % decrease in osteoclast differentiation when compared to the RANKL treated control (p<0.0001). However, mechanical stimulation of the osteoblast was shown to increase the anti-catabolic potential of the secreted EVs, resulting in a significantly more pronounced osteoclast inhibition (37 % reduction, p<0.0001), an effect that is consistent with that seen from the whole secretome (Figure 4 A, B). Osteoclasts differentiated in the presence of osteoblast derived EVs showed decreased number of nuclei when compared to the +RANKL control, indicating a lower level of maturity (Figure 4 A, C). Furthermore, monocyte adhesion wasn’t impaired upon treatment with both static and mechanically activated EVs (Figure 4 A, D). Interestingly, treatment with EVs-depleted CM (DM) led to a notable impairment in monocyte adhesion and a subsequent complete inhibition of osteoclastogenesis. This suggests a preferential packaging of factors within osteoblast EVs that may facilitate monocyte adhesion but also inhibit osteoclast maturation, particularly in response to mechanical stimulation.

**Figure 4.**
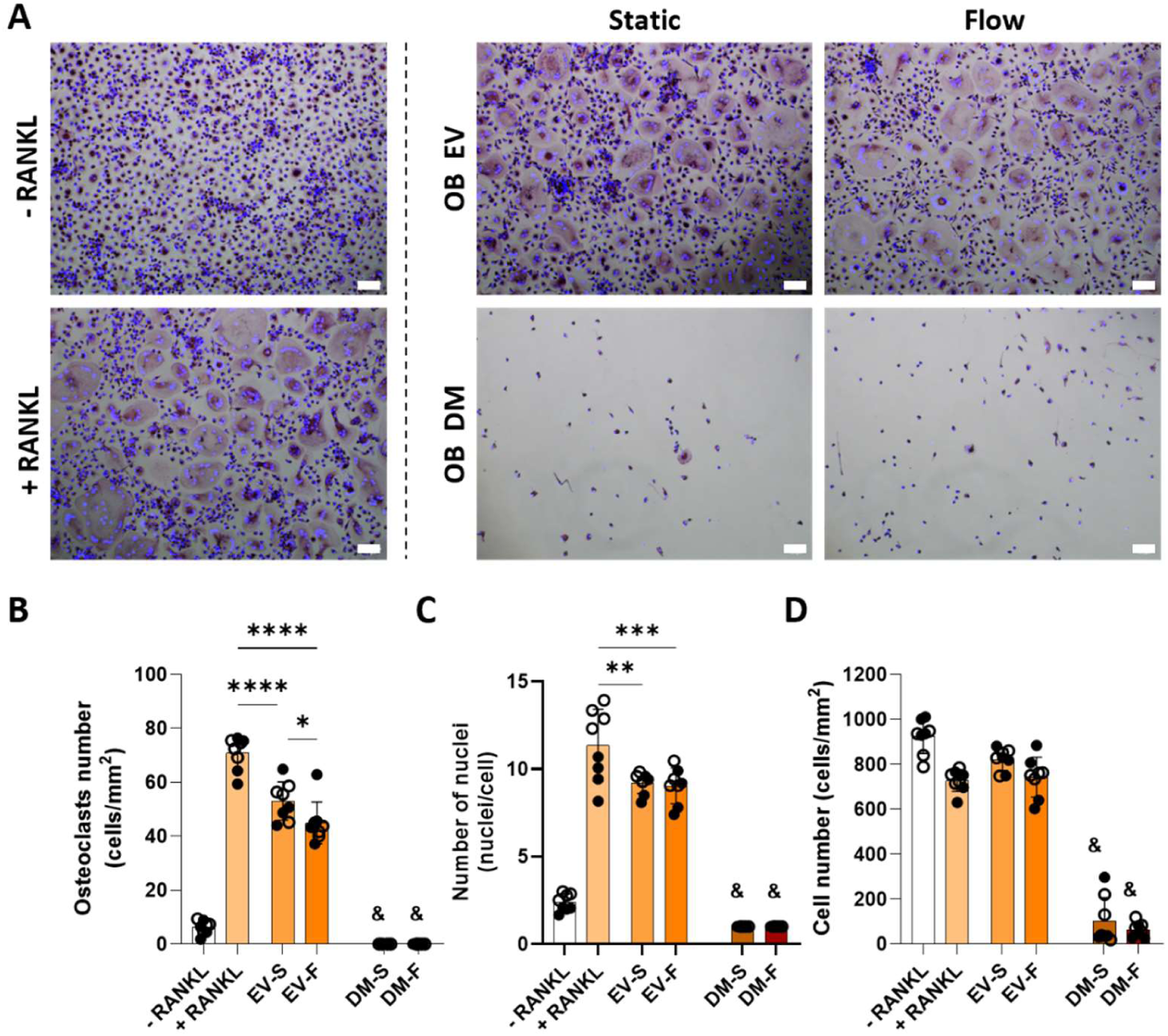
Osteoblasts secrete extracellular vesicles that inhibit osteoclastogenesis, where the inhibitory properties of the EVs are augmented following mechanical stimulation of the osteoblast. **(A)** TRAP and DAPI staining of human monocytes following 7days treatment with M-CSF (-RANKL); M-CSF and RANKL (+RANKL); M-CSF, RANKL and 0.5 µg/mL static osteoblast derived EVs (EV-S); M-CSF, RANKL and 0.5 µg/mL mechanically activated osteoblast derived EVs (EV-F); M-CSF, RANKL and static EVs depleted osteoblast CM (DM-S); and M-CSF, RANKL and mechanically activated EVs depleted osteoblast CM (DM-F). **(B)** Quantification of osteoclast number (identified as TRAP+ cells with ≥3 nuclei), **(C)** nuclei number and **(D)** total cell number. Scale bar 100 µm. Values are mean ± SD for 2 different human donors (● = Donor 1, ○ = Donor 2). *p<0.05, **p<0.01, ***p<0.001, **** p<0.0001, ^&^ p< 0.0001 vs +RANKL, EV-S and EV-F.

Interestingly, similar to that which was observed when treating human monocytes with osteoblast EVs, both static culture and mechanical stimulation of osteocytes resulted in the production of EVs possessing anti-catabolic properties. Quantification of osteoclast numbers revealed that treatment with EVs derived from statically cultured osteocytes resulted in a 54 % reduction in osteoclast formation (p<0.0001). EVs secreted after mechanical stimulation of osteocytes demonstrated a heightened anti-catabolic effect, resulting in a near complete inhibition of osteoclast formation (91 % reduction, p<0.0001) (Figure 5 A, B). In line with what was observed with the other two EVs populations, osteoclasts differentiated in the presence of osteocyte-derived EVs demonstrated a reduced number of nuclei compared to the +RANKL control (Figure 5 A, C). Furthermore, EVs treatment had no impact on the ability of monocytes to adhere. Conversely, CM depleted of EVs (DM) resulted in a substantial impairment of monocyte adhesion, subsequently leading to a lack of osteoclast formation, regardless of mechanical stimulation, in a similar manner to that identified from OB DM. Specifically, the reduction in monocyte adhesion was notably more pronounced when treated with mechanically activated EVs-depleted CM (DM-F) (Figure 5 A, D). This suggests that role of secreted soluble factors in controlling monocyte adhesion and the presence of a selective packaging mechanisms within the EVs that is dependent on the stage of lineage commitment and mechanical environment. The inhibition of osteoclastogenesis elicited by osteocyte derived EVs aligns with that observed upon treatment with the full osteocyte secretome, suggesting that osteocytes coordinate osteoclastogenesis via an EVs-mediated mechanism.

**Figure 5.**
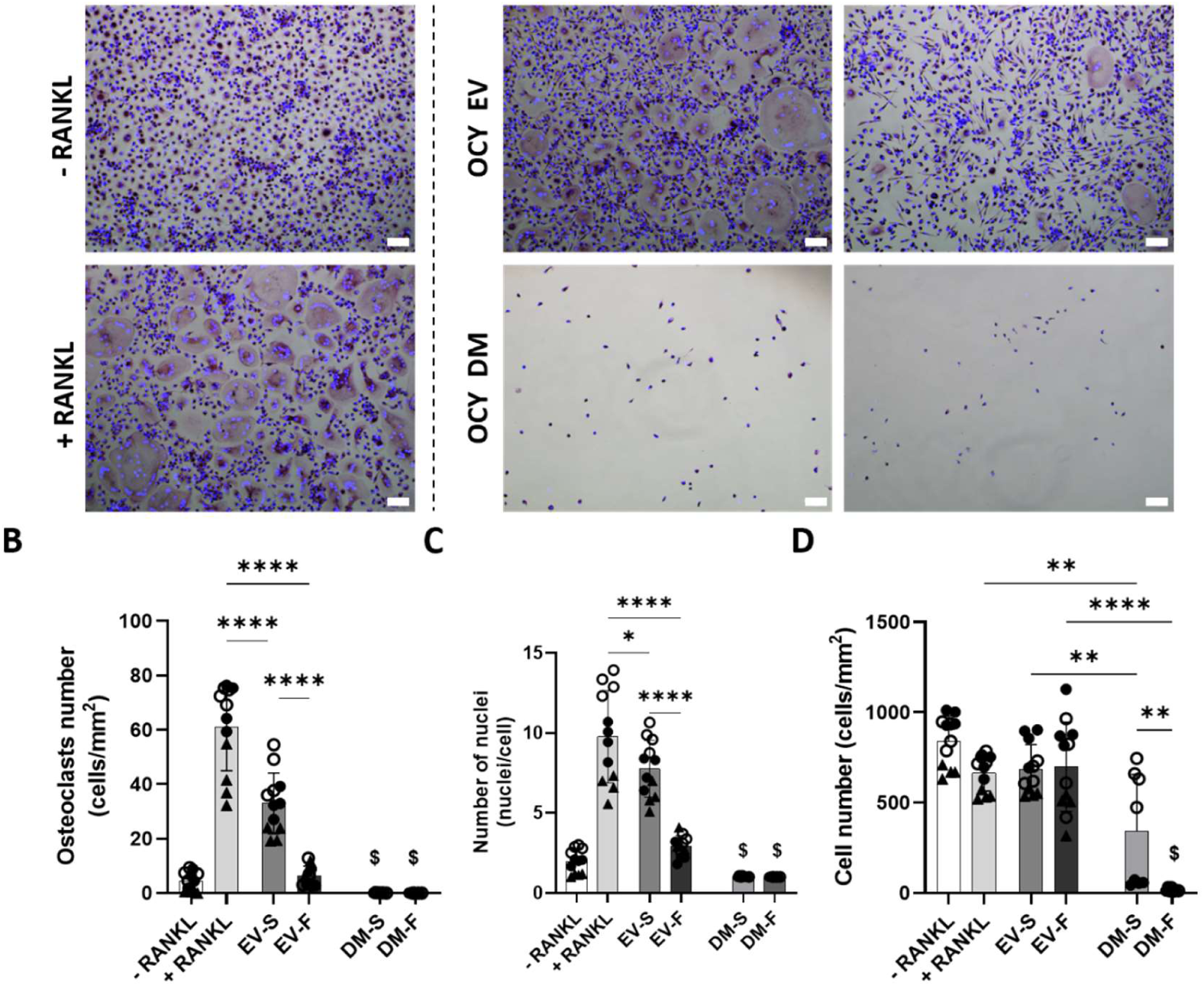
Osteocytes secrete extracellular vesicles that inhibit osteoclastogenesis, where the inhibitory properties of EVs are significantly augmented following mechanical stimulation of the osteocyte. **(A)** TRAP and DAPI staining of human monocytes following 7 days treatment with M-CSF (-RANKL); M-CSF and RANKL (+RANKL); M-CSF, RANKL and 0.5 µg/mL static osteocyte derived EVs (EV-S); M-CSF, RANKL and 0.5 µg/mL mechanically activated osteocyte derived EVs (EV-F); M-CSF, RANKL and static EVs depleted osteocyte CM (DM-S); and M-CSF, RANKL and mechanically activated EVs depleted osteocyte CM (DM-F). (B) Quantification of osteoclast number (identified as TRAP+ cells with ≥3 nuclei), (C) nuclei number and (D) total cell number. Scale bar 100 µm. Values are mean ± SD for 3 different human donors (● = Donor 1, ○ = Donor 2, ▴ = Donor 3). *p<0.05, **p<0.01, ***p<0.001, **** p<0.0001, $ p< 0.0001 vs +RANKL and EV-S.

Taken together, this data demonstrates that mesenchymal derived bone cells can regulate osteoclastogenesis via an EVs-mediated paracrine mechanism that is dependent on the stage of lineage commitment and mechanical environment of the EVs parent cell.

### The anti-bone resorptive properties of the osteocyte derived EVs are strongly dependent on dosage

As the osteocyte is known to be the master orchestrator of bone remodelling, the mechanically activated osteocyte derived EVs possessed the most robust anti-catabolic response, coupled with the fact we have previously demonstrated that these EVs possess both osteogenic and angiogenic properties [35, 36, 45], we next sought to explore the impact of osteocyte-EVs dosage on osteoclastogenesis. In this context we treated human monocytes with increasing concentration of both static and mechanically activated osteocyte derived EVs, namely 0.5, 1, 2 and 4 µg/mL in the presence of both RANKL and M-CSF for 7 days. Interestingly, quantification of osteoclasts revealed that only the treatment with the lower dose of EVs resulted in osteoclast inhibition, mirroring that seen in the CM. However, inhibition was more pronounced in the presence of EVs derived from mechanically stimulated osteocytes. Conversely, doses exceeding the corresponding EVs concentration in the CM - specifically, 1, 2, and 4 µg/mL - prompted an opposite response, enhancing osteoclast differentiation regardless of mechanical stimulation (Figure 6 A, B). In addition, osteoclast nuclei number was upregulated upon treatment with all concentration of EVs, with exception of the lowest EVs dose (0.5 µg/mL), when compared to the +RANKL control group. Mechanical stimulation didn’t affect osteoclast nuclei number following treatment with EVs doses exceeding EVs concentration in the CM, while it elicited a significant reduction following treatment with the lowest EVs dose (Figure 6 A, C). Finally, the total number of cells was not affected by any EVs dose or mechanical stimulation (Figure 6 A, D). These data demonstrate the presence of an optimal dose to maximise the anti-catabolic potential of osteocyte derived EVs.

**Figure 6.**
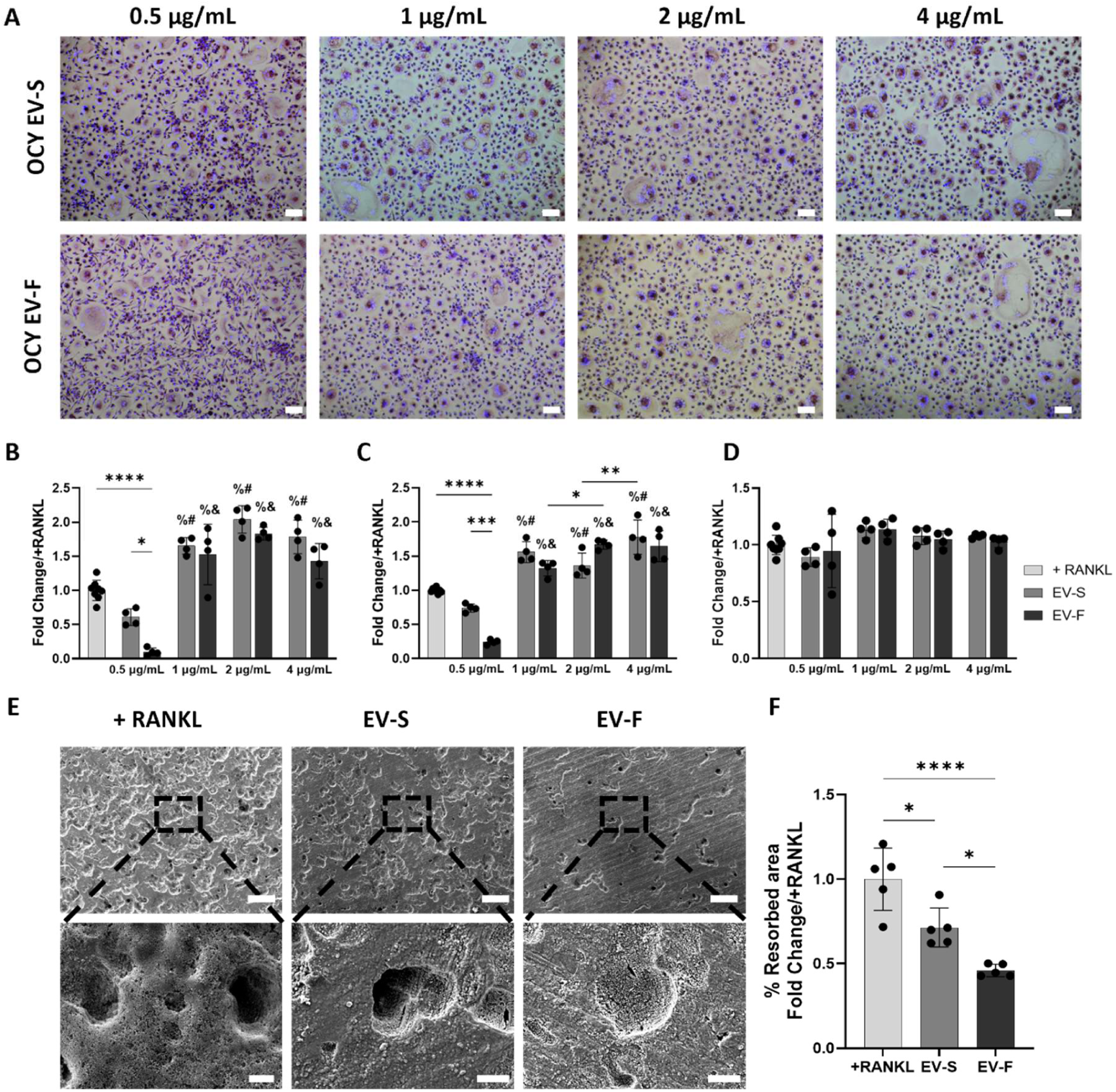
Optimal doses of osteocyte derived EVs inhibit osteoclast differentiation and function. **(A)** TRAP and DAPI staining of human monocytes following 7 days treatment with 0.5, 1, 2 and 4 µg/mL osteocyte-like MLO-Y4 cells derived-static and mechanically activated EVs (Flow) (Scale bar 100 µm). **(B)** Quantification of osteoclast number, (**C)** nuclei number and **(D)** total number of cells. **(E)** SEM images of resorption pits on bovine bone surfaces produced by osteoclasts following OCY-EVs treatment, Scale bars = 200 µm (20 µm-insert) and **(F)** resorbed area quantification. Values are means ± SD. *p<0.05, **p<0.01, ***p<0.001, **** p<0.0001, % p<0.05 vs + RANKL, # p< 0.001 vs 0.5 µg/mL EV-S, & p< 0.0001 vs 0.5 µg/mL EV-F.

**Figure 7.**
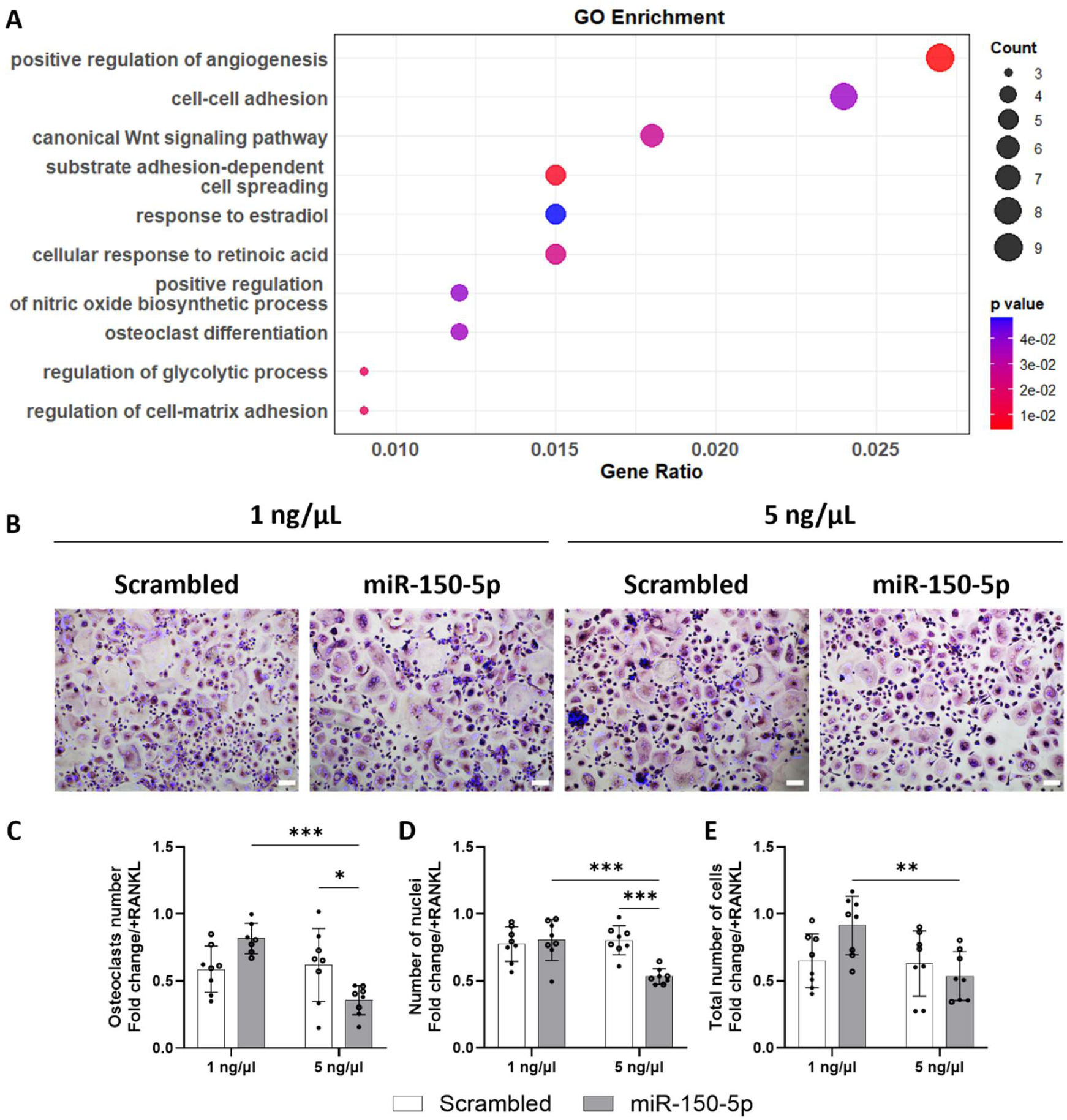
miR-150-5p inhibits osteoclastogenesis in a dose dependent manner. **(A)** GO enrichment analysis of the target genes derived from Target Scan predictions of miR-150-5p. **(B)** TRAP and DAPI staining of human monocytes transfected with 1 ng/µL and 5 ng/µL Scrambled control of miR-150-5p (Scale bar 100 µm). **(C)** Quantification of osteoclast number, **(D)** nuclei number and **(E)** total number of cells. Values are mean ± SD for 2 different human donors (● = Donor 1, ○ = Donor 2). *p<0.05, **p<0.01, ***p<0.001, **** p<0.0001.

After evaluating the anti-osteoclastogenic properties of osteocyte-derived EVs and determining the optimal dose for maximum therapeutic efficacy, we next investigated whether EVs treatment could modulate osteoclast resorptive activity. Treatment with M-CSF and RANKL (+ RANKL control) resulted in resorption of 53 % of the total bone slice area. However, quantification of the osteoclast-resorbed area of bone slices following EVs treatment revealed that EVs derived from both static and mechanically stimulated cultures reduced osteoclast-mediated bone resorption. Notably, EVs secreted in response to osteocyte mechanical stimulation induced a significantly greater reduction in resorbed area compared to both the +RANKL control (62 % reduction, p<0.001) and their static counterparts (48% reduction, p<0.05). Taken together these results showcase osteocyte-derived EVs as promising regulators of both osteoclast differentiation and bone resorptive activity (Figure 6 E, F).

### miR-150-5p is a highly expressed, mechanically regulated miRNA in osteocyte-derived EVs that possesses anti-osteoclastogenic properties

After establishing that osteocytes regulate osteoclastogenesis through an EVs-mediated mechanism, and that EVs derived from mechanically stimulated osteocytes possess enhanced anti-catabolic properties, we proceeded to investigate the specific signalling components within these EVs responsible for the observed effects. In a previous study, we conducted miRNA sequencing on EVs collected from both static and mechanically stimulated osteocytes, identifying miR-150-5p as highly expressed and significantly upregulated in response to mechanical stimulation [36]. To investigate the potential role of miR-150-5p in the anti-osteoclastogenic properties presented earlier, we used Target Scan analysis and identified 356 genes that are potential targets of this mechanically regulated miRNA. Functional enrichment analysis of these genes revealed 112 significantly enriched Gene Ontology (GO) terms. Notably, GO enrichment indicated associations with processes involved in bone regeneration, including “positive regulation of angiogenesis” (GO:0045766) [48], “canonical Wnt signalling pathway” (GO:0060070) [49] and,” positive regulation of nitric oxide biosynthetic process” (GO:0045429) [50]. Additionally, several GO terms were directly related to osteoclast differentiation and function, such as “osteoclast differentiation” (GO:0030316), “regulation of glycolytic process” (GO:0006110) [51], “cell-cell adhesion” (GO:0098609) [52], “regulation of cell-matrix adhesion” (GO:0001952) [53], “cellular response to retinoic acid” (GO:0071300) [54], and “response to estradiol”(GO:0032355) [55] (**Error! Reference source not found.Error! Reference source not found.** A) . Finally, to assess whether miR-150-5p functionally contributes to the anti-catabolic activity of the EVs, human monocytes were transfected with two different concentrations of the miRNA, and osteoclast differentiation was subsequently evaluated. Transfection with the lower dose of miR-150-5p did not significantly affect osteoclast differentiation, multinucleation, or monocyte adhesion compared to the scrambled control. However, transfection with 5 ng/µL of miR-150-5p significantly inhibited osteoclastogenesis, and the resulting osteoclasts displayed fewer nuclei, suggesting impaired maturation. (**Error! Reference source not found.Error! Reference source not found.** B-E). These findings suggest that miR-150-5p, enriched in EVs from mechanically stimulated osteocytes, may play a key role in mediating their anti-catabolic effects by inhibiting osteoclast differentiation, highlighting its therapeutic potential.

## Discussion

Bone is a dynamic tissue that is constantly remodelling via a tightly controlled balance between formation and resorption. This process enables bone to adapt to changing environmental or mechanical cues, adjusting bone mass to maintain an optimal strength to weight ratio. However, when bone resorption exceeds bone formation, it can lead to a net bone loss, microarchitectural deterioration and an increased risk of fracture and/or poor fracture healing. Therefore, this study aimed to characterise how cells from the mesenchymal lineage regulate osteoclastogenesis; how this is influenced by their stage of lineage commitment; and by the dynamic mechanical cues these cells are exposed to physiologically and finally to identify the signalling components mediating this anti-catabolic potential. We demonstrated that the secretome of mesenchymal-derived bone cells inhibited osteoclast differentiation to differing degrees depending on their stage of lineage commitment and on the mechanical environment. Moreover, we have shown that the inhibition of osteoclastogenesis is mediated by EVs released by these bone cells. In addition, we identified the osteocyte as the optimal parent cell for the production of EVs with the most robust anti-catabolic potential and have highlighted the importance of EVs dosage to efficiently regulate both osteoclast differentiation and bone resorptive activity. Finally, we demonstrated that miR-150-5p, cargo of osteocyte-derived EVs, inhibits osteoclast differentiation in a dose dependent manner. Taken together, this study provides an important advancement towards the understanding of EVs-mediated osteoclastogenesis, paving the way for the development of novel nanotherapeutics to prevent bone loss.

Bone cells secrete paracrine factors that inhibit osteoclastogenesis to differing degrees depending on their stage of lineage commitment. We demonstrated that statically cultured MSCs secrete factors that almost completely inhibit osteoclastogenesis, an effect likely mediated by a strong impairment of monocyte adhesion. Conversely, other studies provided evidence of MSCs supporting osteoclastogenesis [9, 56]. These conflicting *in vitro* findings on the impact of MSCs on osteoclastogenesis may stem from variations in the experimental approaches used. Several studies have investigated the interaction between osteoblasts and osteoclasts. In this context, we differentiated osteoblasts from the human MSCs, and our results demonstrated that statically cultured osteoblasts also did not inhibit or promote osteoclastogenesis [57] . These findings demonstrate a change in cell secretome following differentiation along the mesenchymal lineage from the same donor. Osteocytes differentiate from osteoblasts and their role in the regulation of osteoclast activity is well established. Our findings demonstrate that osteocyte secretome has a strong inhibitory effect on osteoclastogenesis. This is supported by previous work demonstrating that osteocyte derived conditioned media dramatically inhibited osteoclast activity, suggesting that osteocytes produce osteoclast inhibitor factors [58, 59]. Taken together, these results demonstrate that bone cells are involved in the regulation of osteoclastogenesis via paracrine signalling. In addition, the differing extents of osteoclastogenesis inhibition indicates that bone cell regulation of osteoclastogenesis is strongly dependent on the stage of lineage commitment.

Bone cells inhibit osteoclastogenesis via a mechanically driven paracrine mechanism. Osteocytes’ ability to respond to mechanical stimulation and regulate osteoclast activity is well known. However, other cells of the osteogenic lineage are also known to be mechanoresponsive, indicating that they might also play a role in mechanically regulated bone remodelling, but the extent to which they can regulate osteoclastogenesis in less well understood. Recent studies demonstrated that mechanical unloading of MSCs promoted osteoclast differentiation [20], while MSCs subjected to oscillatory flow-induced shear stress inhibited osteoclastogenesis [60]. Similarly, our work demonstrated that MSCs subjected to fluid shear stress elicited an inhibition on osteoclastogenesis when compared to the RANKL treated control, but to a lesser extent when compared to the static counterpart. On the other hand, Liu *et al.* observed that both static and dynamic pressure increased osteoclastogenesis [61]. Differences between the studies may be attributed by differences in mechanical regimen and cell species. Osteoblast have also been shown to be involved in the regulation of osteoclastogenesis, and that mechanical cues play a key role in this process [24, 57, 62]. In this context, we demonstrated that osteoblasts subjected to oscillatory fluid flow produced a secretome that inhibited osteoclast differentiation when compared to the static group, in line with previous reports employing other form of mechanical stimulation [63]. Moreover, our results demonstrate that osteocytes subjected to oscillatory fluid flow strongly inhibit osteoclastogenesis via a mechanically driven paracrine mechanism, showing a more pronounced effect than osteoblasts in suppressing osteoclast formation [27]. This is supported by previous studies showing that osteocytes are mechanosensitive, and they inhibit osteoclastogenesis in response to fluid flow [28, 64]. Taken together we have demonstrated the mechanical cues play a key role in bone cell regulation of osteoclastogenesis. When compared to that observed upon treatment with static cultured bone cell conditioned medium, these findings suggest the impact of mechanical stimulation on osteoclastogenesis may differ depending on the mesenchymal lineage stage.

Bone cells from the mesenchymal lineage can coordinate osteoclastogenesis via the release of EVs, a response that can be enhanced following mechanical stimulation. Previous studies have highlighted the pro-anabolic potential of osteoblasts and osteocyte derived EVs and how the anabolic potential can be enhanced by mechanically stimulating the parent cell [35, 65–67]. However, a limited number of studies have investigated whether these EVs can elicit an anti-catabolic response and furthermore delineated the factors involved. Herein, we demonstrate that EVs secreted by both statically cultured and mechanically stimulated MSCs suppressed osteoclastogenesis, an effect mirrored by MSCs-conditioned media (CM) depleted of EVs. This indicates that MSCs secrete both vesicular and soluble inhibitory cues affecting early and late stages of osteoclast development. Previous studies have reported stem cell-derived EVs can suppress osteoclast differentiation and contribute to the regulation of bone resorption under various conditions, including mechanical stimulation [68, 69]. Our data add to this by showing that MSCs-derived EVs reduce osteoclast multinucleation, a hallmark of maturation, without impairing monocyte adhesion, and that their inhibitory effect is not significantly altered by mechanical stimulation. We were also able to demonstrate that by differentiating MSCs into osteoblasts we obtained EVs with a different anti-catabolic potential. Our findings reveal that OB-derived EVs inhibit osteoclastogenesis to differing degrees depending on the OB mechanical environment. EVs from mechanically stimulated OB led to a more pronounced decrease in both the number and maturity of osteoclasts compared to their static counterparts [70]. Interestingly, EVs-depleted OB-CM impaired monocyte adhesion and completely inhibited osteoclastogenesis. Among all cell types, OCY-EVs showed the strongest inhibitory impact on osteoclastogenesis. In particular, mechanically activated OCY-EVs inhibited osteoclast formation by 91 %, aligning with prior evidence that osteocytes are the central regulators of bone resorption during mechanoadaptation [71, 72]. Interestingly, OCY-EVs displayed dose-sensitive behaviour, with low (0.5 µg/mL) doses inhibiting osteoclastogenesis and high doses (>1 µg/mL) promoting it. This biphasic effect suggests a threshold-dependent balance between pro- and anti-catabolic factors, possibly due to changes in cargo saturation or uptake dynamics at higher concentrations [73]. Our findings indicate that optimal dosing is crucial for therapeutic use of osteocyte-derived EVs, especially in pathological states like osteoporosis where remodelling balance is disturbed.

To investigate molecular drivers of the EVs-mediated anti-osteoclastogenic effect, we focused on miR-150-5p, which we previously found to be enriched and mechanically upregulated in osteocyte EVs [36]. GO enrichment analysis highlighted terms related to bone regeneration and osteoclast differentiation and function. Moreover, transfection of miR-150-5p into monocytes resulted in a dose-dependent inhibition of osteoclastogenesis and multinucleation, mirroring the effects of osteocyte EVs. Although miR-150-5p is known to be involved in osteogenesis [74, 75], and monocyte adhesion [76], this study links it to a direct anti-osteoclast activity, expanding its functional repertoire and therapeutic potential. Our data also suggest that the osteocyte EVs-mediated inhibition may be at least partially linked to cargo-specific miRNAs.

One potential limitation of this study is the use of a murine osteocyte cell line to generate OCY-EVs. However, our data demonstrate similar trends in osteoclast formation inhibition and monocyte adhesion following treatments with human OB and murine OCY derived EVs and DM, which differs from the effects observed with human MSCs derived EVs and DM. This suggests that the MLO-Y4 cell line may serve as a reliable model for human osteocyte. In addition, from a therapeutic prospective, previous research has explored the use of plant-derived EVs in human cell treatments, reporting no adverse effects, while supporting human cell proliferation, reducing inflammation and apoptosis [77]. Additionally, reported findings have demonstrated that EVs from human cells can be functionally active and well-tolerated in rodent models [78], suggesting that EVs from different species may represent a safe and viable therapeutic approach for human applications.

In conclusion, this study shows that MSCs, osteoblasts, and osteocytes secrete EVs with the potential to inhibit osteoclastogenesis, although the extent and mechanism of action vary by cell type and mechanical environment. Osteocytes, consistent with their role of master orchestrators of bone remodelling, secrete the most potent EVs, regulating both osteoclast differentiation and function, particularly in response to mechanical cues, a response that is potentially mediated via the mechanical regulation and EVs packaging of miR-150-5p. Combined with the demonstrated anabolic and angiogenic properties of these EVs, mechanically activated osteocyte derived EVs represent a novel multitargeted therapeutic for bone regeneration.

## Acknowledgements

Funding was provided by Research Ireland through the Frontiers for the Future Project Grant (19/FFP/6533) and Award (23/FFP-A/12166) to DAH, by European Union under the Marie Sklodowska-Curie Postdoctoral Fellowship No. 101106209, “METABOLATE” to CG, by EU Horizon 2020 IND, EVPRO to LOD [814495], by the Irish Research Council Advanced Laureate Award EVIC to LOD [IRCLA/2019/49]. The authors gratefully acknowledge the technical support of De. Philippa Timmins and Dr. Pradeep Kumar (Oxford Nanoimaging, UK) with super resolution microscopy.

## Data availability statement

The data are available from the corresponding author on reasonable request

## Declaration of Interests

The authors declare no conflict of interest.

## Author contributions

**M.M.:** study design, investigation, data analysis, visualisation, writing – original draft. **C.M.:** contribution to investigation. **R.A.: e**xtracellular vesicles collection and characterisation. **S.P.:** contribution to investigation. **M.Y.B.:** contribution to investigation, writing – review and editing. **L.A.M.:** SEM samples preparation and imaging. **C.G**.: extracellular vesicles collection. **T. NN.:** contribution to investigation. **C.T.B:** contribution to study design, technical support, interpretation of data, **L.O.D.:** contribution to study design, technical support, interpretation of data**. D.A.H.:** conceptualisation, study design, resources, writing – original draft and review and editing, supervision, project administration and funding acquisition. All authors critically revised the manuscript and approved the final version.

## Notes

### Competing Interest Statement

The authors have declared no competing interest.

